## Supplementary File 1 for "Multi-omic temporal landscape of plasma and synovial fluid-derived extracellular vesicles using an experimental model of equine osteoarthritis"

**Supplementary File 1.** Clustering analysis to determine the effect of missing values.

A. Cluster analysis of plasma-derived EVs, and B. SF-derived EVs, C. Density plot of plasma-derived EVs showing that when all the rows with missing values were removed the overall distribution shifted higher, D, Density plot of SF-derived EVs; the maximum number of proteins consistently identified in the samples was 50 and that was still in only 26 of the samples. Many proteins were identified in <10 of the samples, which may be indictive of the SF being lower quality SF samples might be generally variable.

A. Cluster plot for plasma-derived EVs

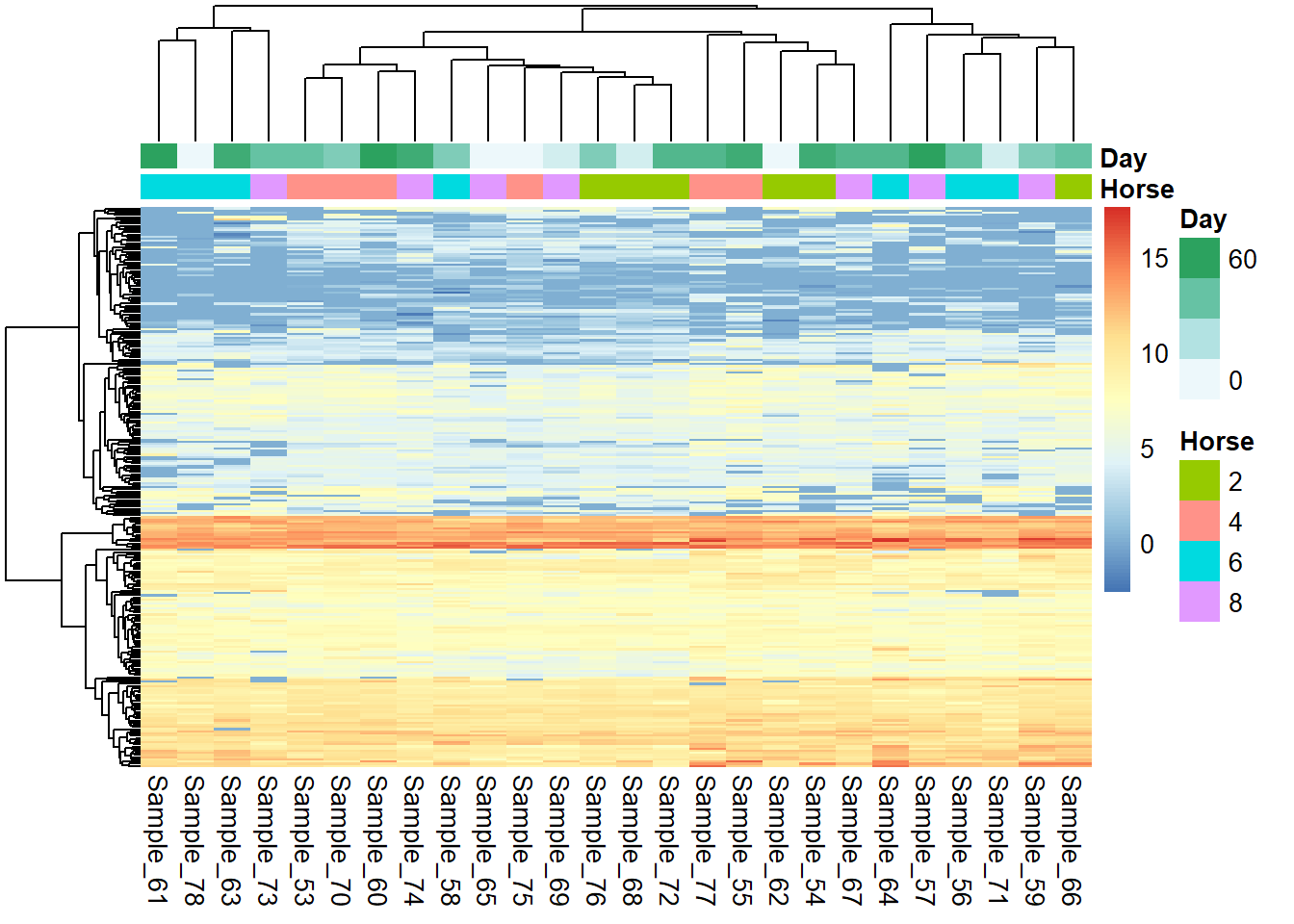

B. Cluster plot for SF-derived EVs

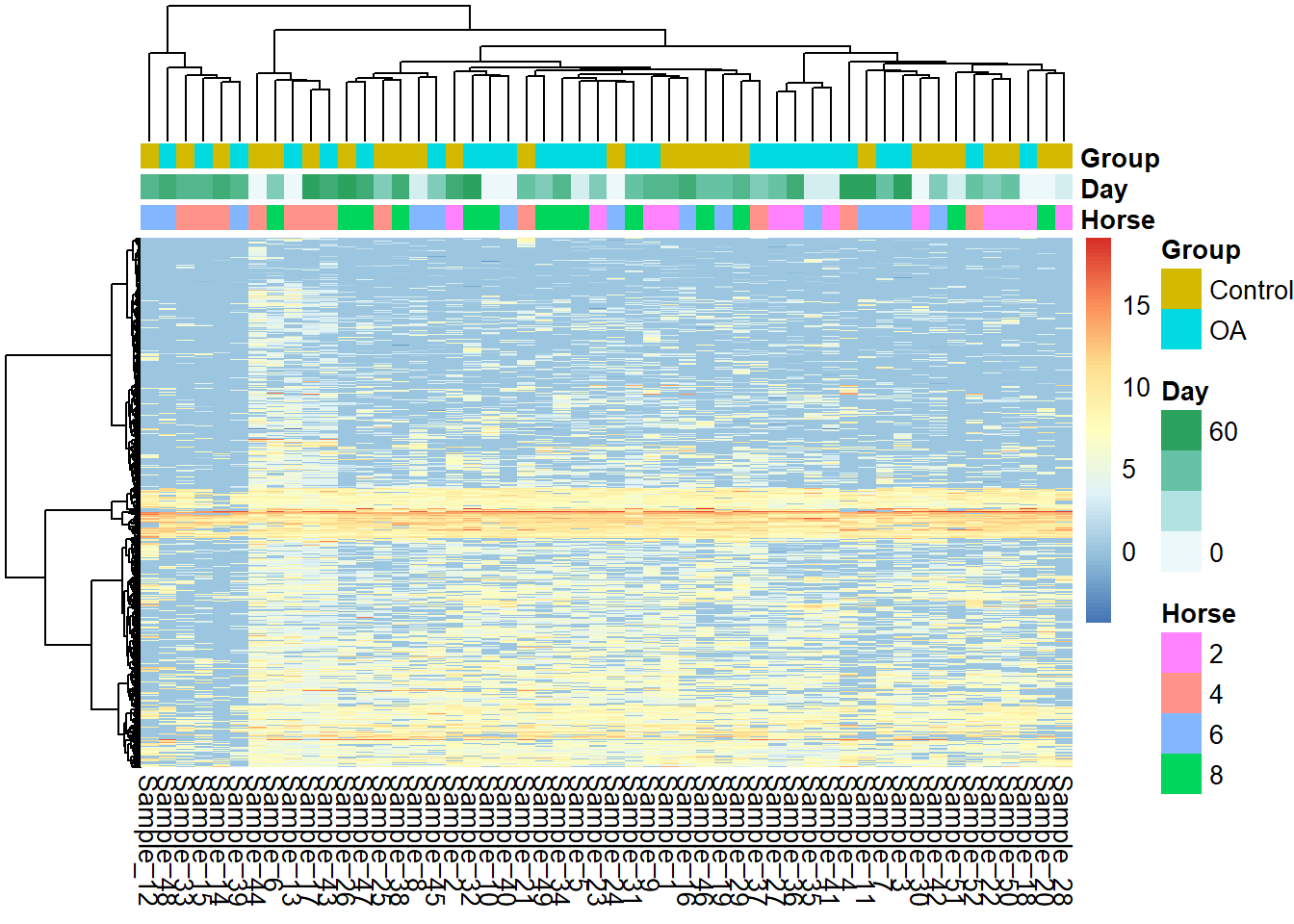

C. Density plot for plasma-derived EVs

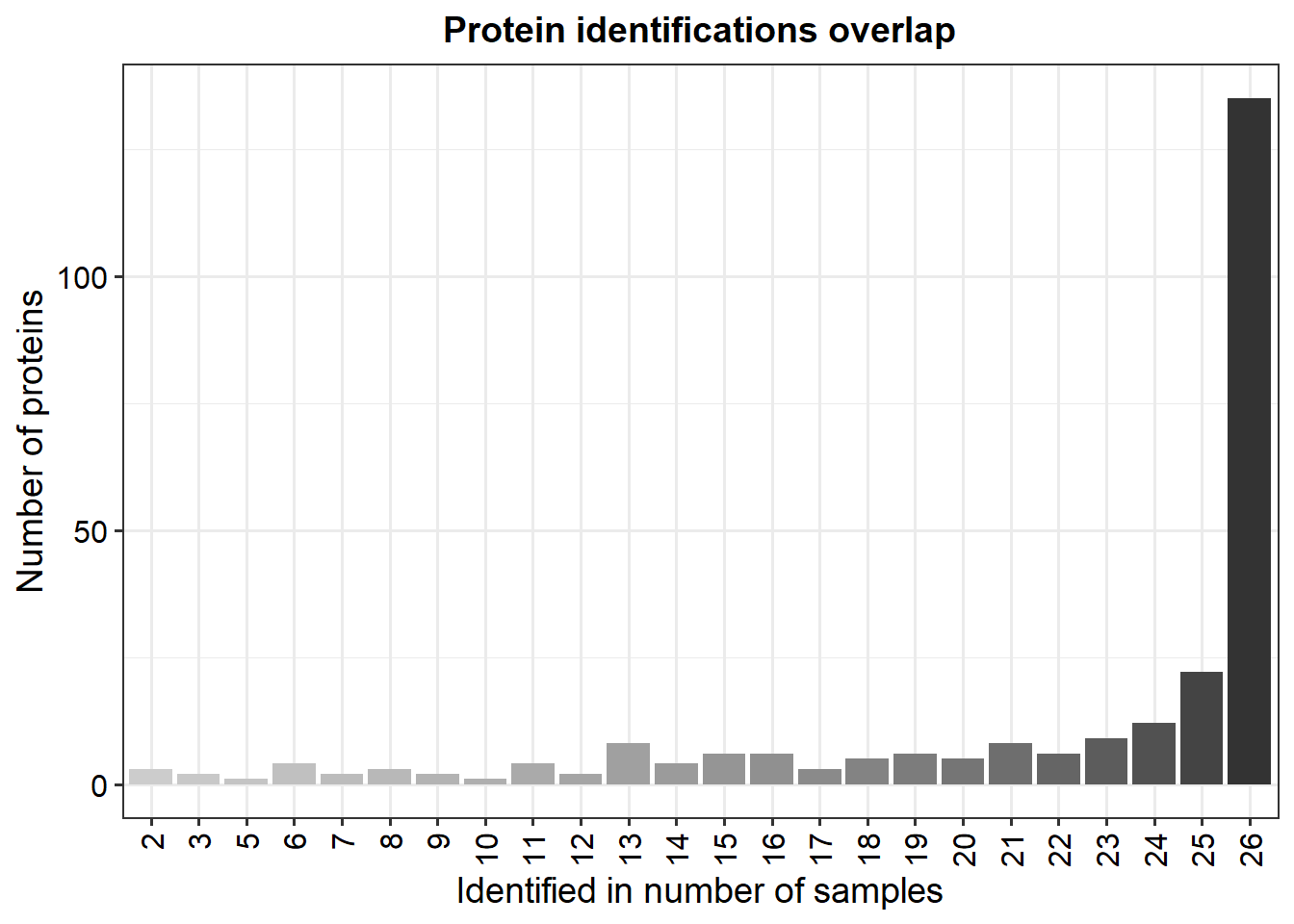

D. Density plot for SF-derived EVs

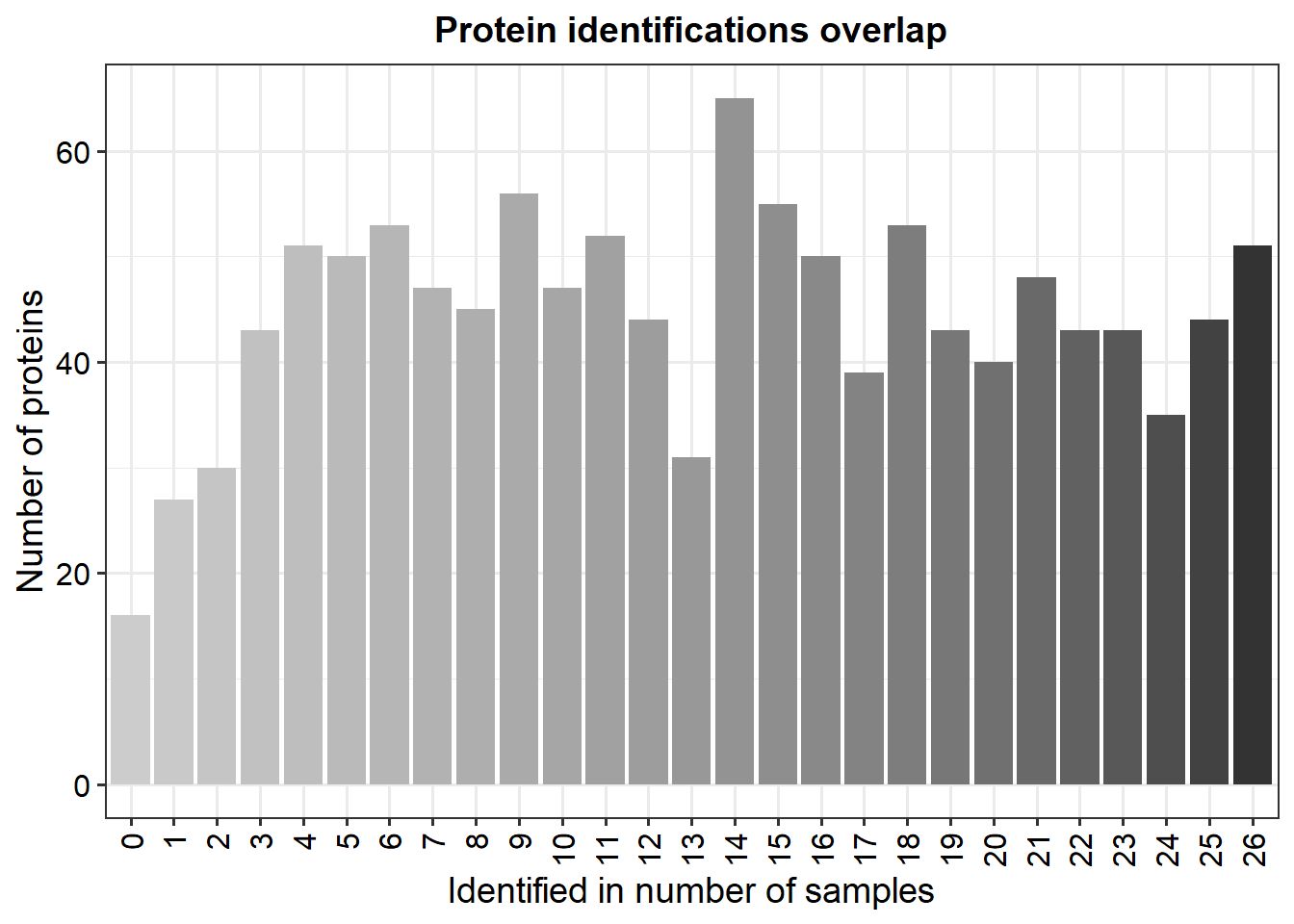
