## Supplementary File 3 for "Multi-omic temporal landscape of plasma and synovial fluid-derived extracellular vesicles using an experimental model of equine osteoarthritis"

Supplementary File 1. Assessment of whether the missing values in small RNA sequencing data were true zeroes or sampling zeroes using Pearson’s correlation of total number of missing values by sequencing depth for A. miRNA plasma EVs, B. lncRNA plasma EVs, C. snoRNA plasma EVs, D. snRNA plasma EVs, E. miRNA SF EVs, F. lncRNA SF EVs, G. snoRNA SF EVs and H. snRNA SF EVs.

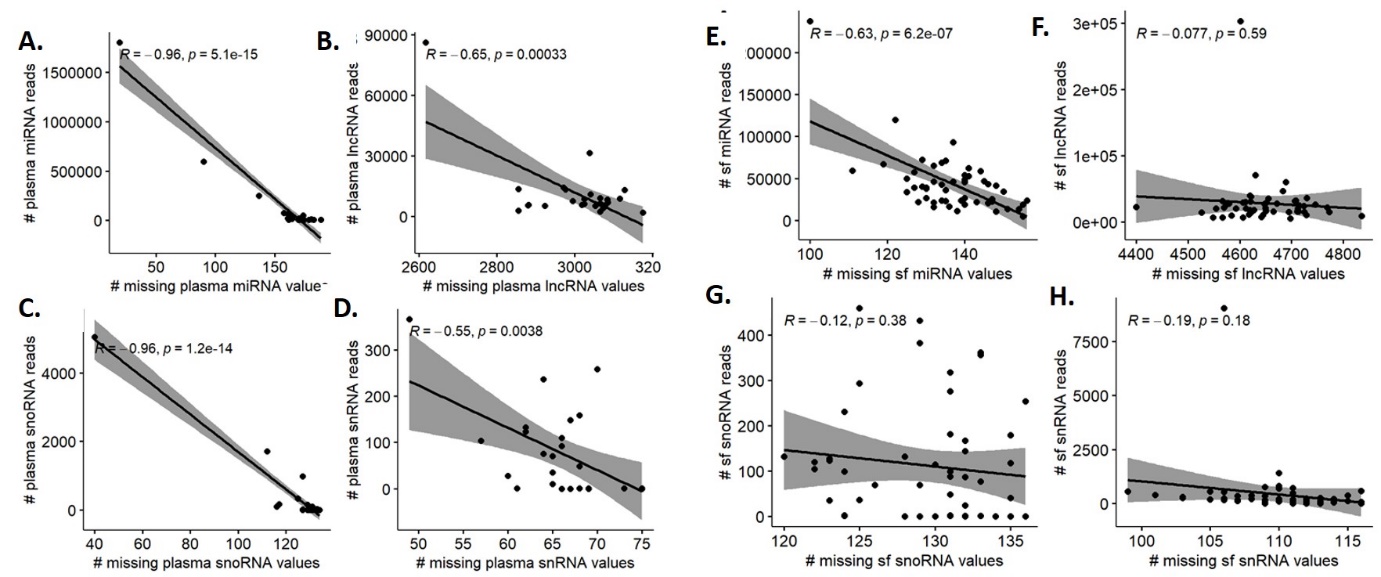
