## Supplementary File 4 for "Multi-omic temporal landscape of plasma and synovial fluid-derived extracellular vesicles using an experimental model of equine osteoarthritis"

Supplementary File 2. Assessment of whether the missing values in small RNA sequencing data were true zeroes or sampling zeroes using PCA with complete observations compared to PCA with missing values as zeroes. A. Without missing values, B. Missing values as zeros, C. Correlation map without missing values, D. Correlation map with missing values. When the missing values were included as 0’s PC1 had an almost 1:1 correlation with sequencing depth. Example shown is for miRNAs.

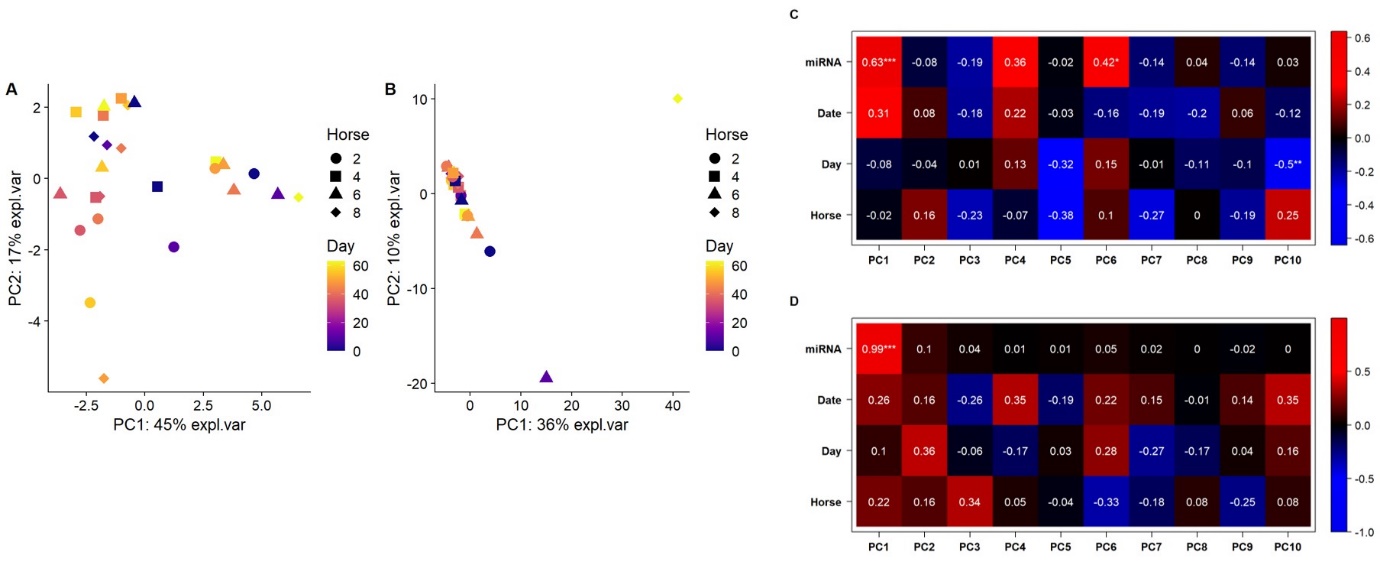
